# Cryopreservation effect on DNA methylation profile in rainbow trout spermatozoa

**DOI:** 10.1101/2023.08.11.552935

**Authors:** Marina El Kamouh, Aurélien Brionne, Audrey Laurent, Catherine Labbé

## Abstract

Spermatozoa are the cells the most commonly used for cryopreservation of valuable genetic resources in aquaculture. It is known that fish spermatozoa transmit to the embryo not only their genetic but also their epigenetic profile, particularly DNA methylation. Therefore, any alteration of the DNA methylation profile in spermatozoa induces the risk of transmitting epigenetic alterations to the offspring. The aim of this study was to assess the effect of cryopreservation on DNA methylation in rainbow trout spermatozoa. To trigger variable cellular response after freezing-thawing, spermatozoa from mature males were cryopreserved with dimethyl sulfoxide, methanol or glycerol as cryoprotectant. We observed that dimethyl sulfoxide was the best to preserve thawed spermatozoa functions. Methanol preserved only slightly all the cellular parameters, while glycerol failed to protect motility and fertilization ability. The consequences on DNA methylation were assessed using Reduced Representation Bisulfite Sequencing (RRBS). Sperm cryopreservation did not thoroughly impact DNA methylation, although 335 to 564 differentially methylated cytosines were characterized depending on the cryoprotectant. Very few of them were shared between cryoprotectants, and no correlation with the extent of cellular damage was found. Our study showed that DNA methylation was only slightly altered after sperm cryopreservation, and this may render further analysis of the risk for the progeny very challenging.

## INTRODUCTION

Cryopreservation of fish spermatozoa is widely used to secure genetic selection programs and to preserve rare or endangered species. It is known that besides conveying the genetic background of the breeders to the progeny, the fish spermatozoa also bear a specific epigenetic profile, particularly DNA methylation. This profile is important for the offspring as it has been shown in zebrafish that some features of the sperm methylome are inherited and maintained during early embryogenesis, and that they contribute to the regulation of zebrafish embryo development ^1–3^. Such sperm methylome inheritance by the embryos raises the concern that any alteration of DNA methylation in spermatozoa bears the risk of transmitting epigenetic alterations to the offspring. The effect of cryopreservation on sperm DNA methylation has been little explored yet, and the studies are reduced to few species, using very different methylation analysis methods. In human, two candidate analysis of parentally imprinted genes showed that sperm cryopreservation had no effect on the maternally imprinted genes LIT1, SNRPN, MEST and SNURF-SNRPN, on the paternally ones MEG3, H19, and UBE3A, nor on repetitive elements (ALU, LINE1), on a spermatogenesis-specific gene (VASA) or on a gene associated with male infertility (MTHFR) ^4,5^. However, a third candidate gene study showed more recently that sperm cryopreservation induced an increased methylation in the promoter regions of three other paternally imprinted genes (PAX8, PEG3 and RTL1) ^6^. More global analyses in ram ^7^ and stallion ^8^ using ELISA showed an increased DNA methylation in frozen-thawed spermatozoa, but the affected genomic regions were not explored. In fish, some studies also evaluated the global DNA methylation of cryopreserved spermatozoa, using LUMA (Luminometric assay) which provides information on the overall DNA methylation, without providing information on the identity of the altered sites ^9,10^. These studies showed intricate species-related effects and cryoprotectant-related ones. Indeed, methanol (MeOH) did not alter DNA methylation in European eel and in goldfish, but it triggered a slight hypermethylation in zebrafish. Goldfish DNA methylation was not completely insensitive to cryopreservation since dimethyl sulfoxide (DMSO) and 1,2-propanediol did induce some DNA hypomethylation in this species ^10^. Cryopreservation also induced a global hypomethylation (assessed with LUMA) of tambaqui spermatozoa whatever the tested cryoprotectants, including MeOH, DMSO and glycerol ^11^. In all, cryopreservation might induce a risk of DNA methylation alteration in spermatozoa, but the extent of the damages at the scale of the genome and its functional regions has still not been addressed.

Rainbow trout (*Oncorhynchus mykiss*) is important in aquaculture and it has been an ever-present species in selective breeding programs in the past decades ^12,13^. Cryobanking of the improved commercial strains using cryopreserved sperm is operational in France ^14^, but to date, the impact of sperm cryopreservation on DNA methylation has never been studied in rainbow trout. The aim of our study was to assess whether cryopreservation of rainbow trout spermatozoa can alter their DNA methylation profile and whether alterations of the cellular quality of thawed sperm is related to possible DNA methylation alterations. To this end, we used different cryoprotectant molecules in order to induce some variability in cellular cryoprotection, namely DMSO, MeOH and glycerol. DMSO and MeOH are the most commonly used molecules for sperm cryopreservation in salmonids. However, DMSO is a highly chemically reactive molecule comparing to MeOH. Glycerol does not protect spermatozoa functionality as do DMSO and MeOH, but it still provides some structural protection. Thawed spermatozoa quality was evaluated from their membrane quality, mitochondria quality, motility and fertilization ability. DNA methylation was assessed at the genome-wide level using reduced representation bisulfite sequencing (RRBS), a method that allows a partial genome representation, thereby allowing the assessment of many biological replicates for each cryoprotectant.

## RESULTS

### Cellular quality of the thawed spermatozoa according to the cryoprotectant

Plasma membrane quality of the thawed spermatozoa was deduced from spermatozoa ability to exclude the membrane impermeant propidium iodide. As shown Figure 1a, DMSO preserved particularly well the plasma membrane, since all samples had more than 75 % of the cells with intact plasma membrane. Glycerol was also able to preserve the plasma membrane quality with an efficiency similar to that of DMSO. On the contrary, MeOH yielded the lowest percentages of cells with intact plasma membranes, and the highest variability in plasma membrane quality among the spermatozoa of the different males. Mitochondria quality of the spermatozoa was estimated from their JC-1 staining intensity (Figure 1b). Again, DMSO and glycerol showed the best ability to preserve the mitochondria quality, with more than 50 % of the cells having mitochondria with a high activity, while the spermatozoa cryopreserved with MeOH had a very low mitochondria quality.

**Figure 1:**
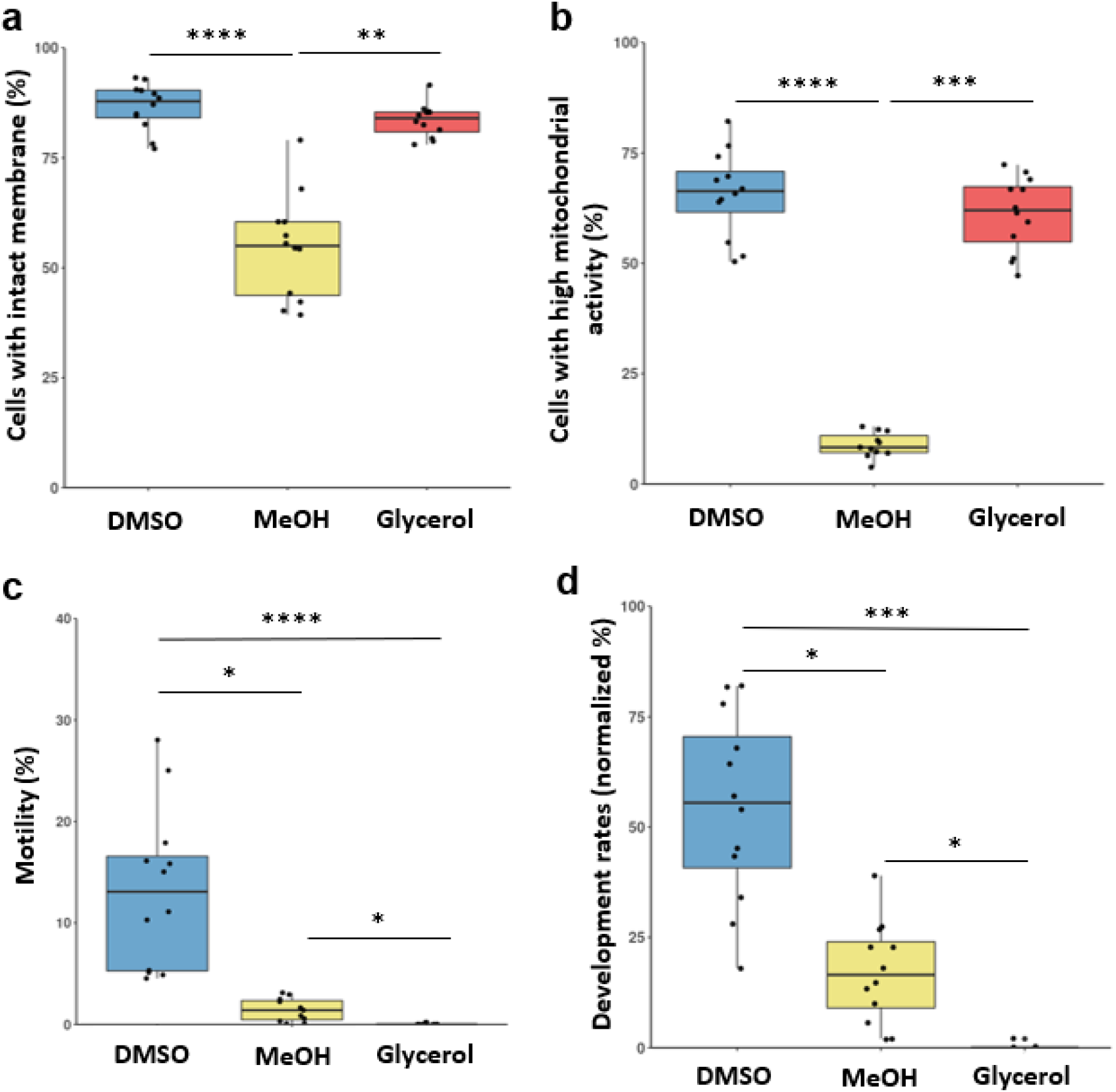
Cellular quality of the thawed spermatozoa according to the cryoprotectant. a. Cells with an intact plasma membrane, analyzed from the percentage of cells which excluded the impermeant fluorochrome propidium iodide; b. Cells with high mitochondrial activity, analyzed from the percentage of cells which had the highest fluorescence intensity after JC-1 labelling; c. Motile cells calculated from the percentage of cells which had an average path velocity ≥ 30 µm/s; d. Development at eyed stage after fertilization with cryopreserved spermatozoa. The percentages were normalized according to the development rate of the egg quality controls (fertilized with standard fresh sperm). In the boxplot representation (a to d), each dot corresponds to individual value for one sperm (n=12 biological replicates in each condition). The boxplots show (from the bottom to the top): minimum, 25% percentile, median, 75% percentile and maximum. Significant differences between cryoprotectants (Kruskall & Wallis test) are indicated by *: p value < 0,05; **: p value < 0,01; ***: p value < 0,001,****: p value < 0,0001.

Regarding motility, despite the overall low sperm motility after thawing (Figure 1c), the highest values were obtained with DMSO. The spermatozoa cryopreserved in MeOH retained very little motility while glycerol completely fail to protect sperm motility ability. The same trend was observed with the fertilization ability of the cryopreserved spermatozoa (Figure 1d). Fertilization is a test that comprehensively reflects the integral functionality of the spermatozoa. Our results showed that the best fertilization ability was obtained with DMSO. MeOH yielded lower performances, with no fertilization rates above 50 %. After cryopreservation in glycerol, the spermatozoa completely lost their fertilization ability, which was to be expected when considering the total loss of motility of these samples. These fertilization results were confirmed in another fertilization test in which other straws from the same sperm samples were used, with another batch of oocytes (results not shown).

In all, cryopreservation with different cryoprotectants altered the functionality of the thawed spermatozoa, and the different cryoprotectants yielded different cellular response to cryopreservation. We observed that DMSO preserved particularly well the membrane quality and the mitochondria quality, and protected to some extent the motility and the fertilization ability. MeOH was less efficient at protecting whatever cellular quality level, while interestingly, glycerol was able to preserve well the membrane quality and mitochondria quality, but was completely inefficient regarding the motility or the fertilization ability. This wide set of alterations depending on the samples provided us with a very variable biological material to explore further the risks of changes in DNA methylation after cryopreservation.

### Validation of the RRBS sequencing data for DNA methylation analysis

Because RRBS yields a reduced representation of the analyzed genomes, we assessed whether our sequencing data still had enough cytosines in CpG sites to provide for an acceptable representation of those in the whole rainbow trout genome. As a reference, we used a total number of cytosines (in CpG sites) of 70.6 × 10^6^, although the reference trout genome contains 35.3 × 10^6^ CpG dinucleotides (Omyk_1.0 genome version). Indeed, cytosines of both forward and reverse strands can be sequenced (named plus and minus strands depending on their sense on the reference genome), and this can result in the sequencing of the same CpG site twice. We did not remove this redundancy (+ and – CpGs) from our data, because we do not know whether a CpG site can be altered symmetrically by cryopreservation or not. As a consequence, the whole information was kept, and each sequenced cytosine (in a CpG site) in our conditions is one out of the 70.6 × 10^6^ genomic cytosines (twice the number of genomic CpG sites). We found an average of 2.9 × 10^6^ cytosines in CpG sites per sample (fresh and cryopreserved sperm), that represent 4 % of the 70.6 × 10^6^ genomic cytosines in CpG sites (Table1). This representation was slightly higher in MeOH, likely because these samples also had more reads than those in the other conditions, leading to more successful extraction of the cytosines. It is noteworthy that when combining all the cytosines of all the males from one condition, the representation of the genomic cytosines in CpG sites was doubled (9 %). Hence, the reduced representation of the genomic cytosines in CpG sites by the data at the sample level was compensated for by the large number of biological replicates in this study.

To investigate whether these 9 % of genomic cytosines in CpG sites were homogeneously distributed over the entire reference genome, we analyzed their individual location. We observed a homogeneous distribution of the cytosines all along the chromosomes of the rainbow trout genome, with never more than 150 kb between two consecutive cytosines (Figure 2). The same homogeneous distribution of the sequenced cytosines in CpGs sites was found within each sample of each cryopreservation condition. Therefore, this distribution indicates that the captured cytosines in CpG sites provided a reduced but homogeneously distributed representation of the rainbow trout reference genome.

**Figure 2:**
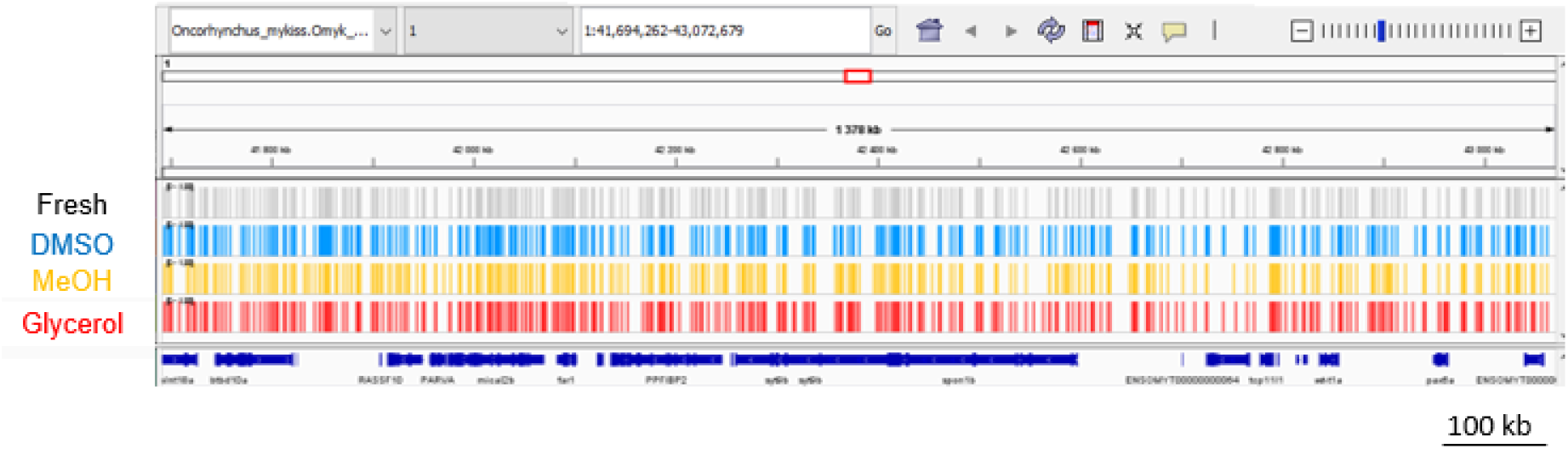
Example of the homogeneous distribution of the sequenced cytosines in CpG sites over the reference genome. The displayed region is representative of the pattern observed with all the sequenced data over the whole genome. The chart was obtained with IGV_2.8.13 and shows the distribution on the chromosome 1 from the position 58 234 274 to 59 476 270 (1.2 × 10^6^ bases) of the rainbow trout reference genome “Omyk_1.0 genome version”. Each line corresponds to a different condition in which all the sequenced cytosines (in CpG sites) of all 12 males are represented. Each sequenced cytosine in a CpG site is represented by a bar. The bottom line represents the genes (dark blue) associated to this genomic fraction.

### Global DNA methylation after cryopreservation

Cryopreservation had little effect on the sperm global DNA methylation. Indeed, we found that the average DNA methylation percentage of cryopreserved sperm (DMSO: 86.60 ± 1.05, MeOH: 87.34 ± 0.11 and glycerol: 85.93 ± 2.06) were not significantly different (p>0.05, Kruskal-Wallis test) from the fresh control one (86.34 ± 1.76). This indicates that cryopreservation did not impact DNA methylation at a global scale. In a further analysis of the data, instead of using only the average methylation ratio of each sample, we compared all the samples to each other based on the methylation ratio of every sequenced cytosine (PCA, Figure 3a). We found no clear segregation between fresh and cryopreserved sperm regardless of the cryoprotectant used. Only four samples were distant from the others to some extent (on the first axis of the PCA), but they did not share any treatment similarities (2 fresh samples, one DMSO, and one glycerol). Their separation from the others can be explained by the lower number of cytosines in CpG sites of these specific samples. This result enforces the fact that cryopreservation did not thoroughly affect DNA methylation, and that if any alteration has been induced by cryopreservation, it should be sought at a more discrete level.

**Figure 3:**
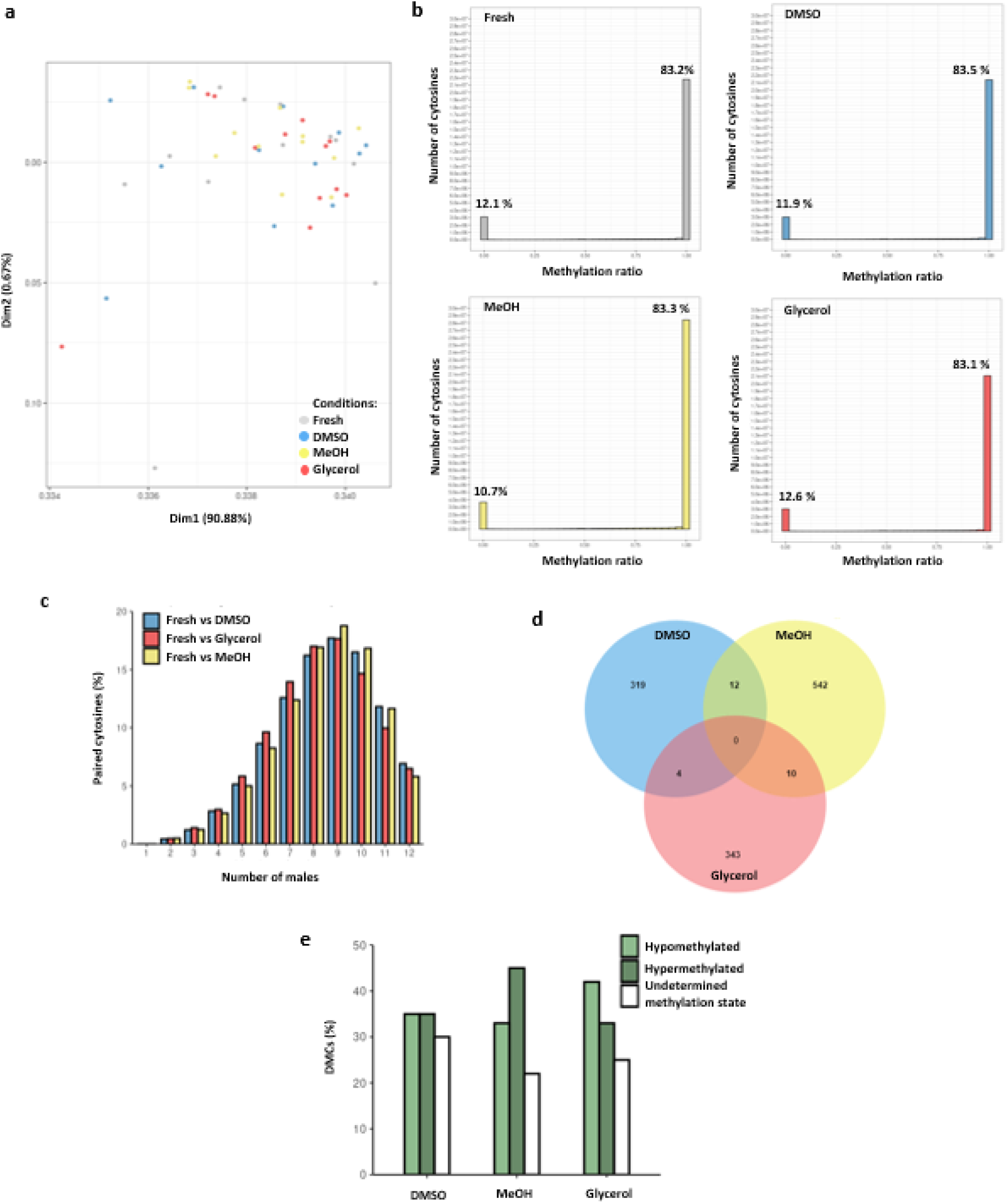
Analysis of sperm DNA methylation according to the cryoprotectant. a. Principal Component Analysis showing the distribution of the fresh and cryopreserved sperm samples according to the DNA methylation ratio of each cytosine within each individual sample (n= 12 sperm samples per condition). No clear segregation was observed among the different treatments. b. Frequency plot showing the number of cytosines in CpG sites according to their methylation ratio (0 to 1) for each condition (fresh and cryopreserved with DMSO, MeOH and glycerol). The x axis shows 20 classes of 0.05 interval values (methylation ratios ranging from 0-0.05 to 0.95-1.0). The percentages written above the classes 0-0.05 and 0.95-1.0 indicate the percentage of the total cytosines in CpG sites in these 2 classes. c. Histogram showing the distribution of the number of paired cytosines according to the number of males sharing them, in each comparison between fresh and cryopreservation with a given cryoprotectant. Results are expressed as a percentage of the total number of paired cytosines (fresh vs DMSO: 748 770; fresh vs MeOH: 1 645 797; fresh vs glycerol: 714 554). d. Venn Diagram of differentially methylated cytosines (DMCs) obtained in each comparison between fresh and each cryoprotectant. The numbers indicate the number of DMCs in each group. e. Histogram showing the distribution of the different classes of DMCs in each comparison: hypomethylated (>50 % of males having this DMC as hypomethylated), hypermethylated (>50 % of males having this DMC as hypermethylated) and unclassified methylation status (DMCs whose methylation status between males was too variable to be included into one of the first two classes). Values are expressed as a percentage of total DMCs: (335 in DMSO, 564 in MeOH and 357 in glycerol).

We thus studied the individual methylation ratio of all the cytosines with a sequencing depth N ≥ 10. For all treatments, more than 80 % of the analyzed cytosines in CpG sites had a methylation ratio of 1 (Figure 3b, class 0.95-1). This confirms the highly methylated status of the cytosines in all samples, as shown above in the global results. The second most represented category encompassed the cytosines with a methylation ratio of 0. Only a very small proportion of cytosines had intermediate methylation values, meaning that there is very little variability in the methylation status of the cytosines between spermatozoa within a sample (most are in the binary status of either methylated or unmethylated). When we compared the fresh to the cryopreserved samples, we did not observe any shift between the 3 categories (Figure 3b, percentages) that would have indicated that some cytosines would have acquired a more variable status among spermatozoa. Besides, no significant difference was found when comparing the 12 fresh samples to any of the cryopreserved ones (p>0.05, Kruskal-Wallis test). Thus, if sperm cryopreservation has induced some subtle alterations of the cytosine methylation ratio, it was not detected from this representation.

### Characterization of differentially methylated cytosines (DMCs) after sperm cryopreservation

The differential methylation of cytosines in CpG sites between fresh and cryopreserved samples was characterized using a paired-driven analysis, in order to focus on the cytosines within each male that were present in both the fresh and respectively the DMSO, MeOH and glycerol conditions. This prevented any statistical bias where neighboring cytosines are used in the comparison to compensate for the lack of information at a given site. These cytosines will be referred to as paired cytosines (i.e. a cytosine at a genomic position that is present in both the fresh sperm and the corresponding cryopreserved one). In this analysis, all cytosines with a sequencing depth N ≥ 1 were included. We first assessed how many paired cytosines would be shared between the different males. Surprisingly, the highest proportion of these cytosines (> 15 %) were shared by 8 to 10 males (Figure 3c) whereas fewer cytosines (< 6 %) were present in only 1 to 5 males. Furthermore, the number of paired cytosines shared from 6 to 12 males represented 90 % of the total number of paired cytosines. That paired cytosines were found as being more abundant in more males would indicate that some fragments were more prone to be sequenced than others in the majority of the males, although the origin of this phenomenon regarding a possible sequencing bias is difficult to understand. In all, since we wanted to select the highest proportion of paired cytosines, we selected the paired cytosines present in at least 6 males (n ≥ 6). This threshold also allowed us to strengthen the biological meaning of the DMCs that we may obtain.

When comparing DMSO, MeOH and glycerol to fresh sperm, we found respectively 335, 564 and 357 DMCs (Figure 3d). No DMCs were common to the three cryopreservation conditions, and only few DMCs were common to 2 cryoprotectants. None of these DMCs had a straightforward status as we observed that depending on the males, a given DMC could be hyper-or hypomethylated after cryopreservation. In order to sort out some trends in the methylation changes, the DMCs were classified as hypermethylated, or hypomethylated, when they had the same trend in > 50 % of the males. As shown in Figure 3e, cryopreservation with DMSO triggered the same number of hypomethylated and hypermethylated DMCs (35 %) whereas in MeOH, more DMCs were hypomethylated (45 %) than hypermethylated (33 %). A reverse tendency was observed after cryopreservation with glycerol where more DMCs were hypermethylated (42 %) than hypomethylated (32 %). Some DMCs remained unclassified because they did not show a clear trend among the corresponding males. This emphasizes the fact that cryopreservation did not trigger a homogeneous effect in all the males, and that sperm cryopreservation impact on DNA methylation was different depending on the cryoprotectant.

### Characterization of regions potentially sensitive to cryopreservation

Despite the low DMCs numbers obtained in our cryopreservation conditions, we investigated whether there were any DMR (differentially methylated region) between fresh and each cryoprotectant. No DMRs were found after cryopreservation with DMSO and MeOH, whereas only 2 DMRs were observed after cryopreservation with glycerol. Some potentially sensitive regions could have escaped detection by DSS because of the stringency of the chosen settings, where a DMR search requires that at least five consecutive CpGs are present in a sliding 50 bases window. In an empirical approach focused on the DMCs that we have identified, we sought for regions with at least two DMCs separated by no more than 100 bp (our maximum read size). These regions were referred to as potentially sensitive regions. We found 62 of these potentially sensitive regions, present in at least one comparison between cryopreserved sperm and fresh controls (Table 2). Among them, 17 were sensitive to DMSO, i.e. they contained at least 2 DMCs in the DMSO-fresh comparison, 33 were sensitive to MeOH, and 16 were sensitive to glycerol. One region was affected by all 3 cryoprotectants (8 : 79 446 507 - 8 : 79 446 591, table 2), and 3 regions were sensitive to both DMSO and MeOH, but not to glycerol.

These regions found to be sensitive to the cryoprotectants were distributed in all genomic regions and these included features such as introns, intron1, exons, promoter 5kb and intergenic regions (Table 2). Some annotated genes were identified in these regions, although the only region common to the 3 cryoprotectants belonged to an intergenic region. It is noteworthy that among the regions related to annotated genes (promoters, introns and exons), only 4 of them were affected by DMSO while 12 were affected by MeOH and 12 by glycerol. We found that the distal promoters of *ebf2* and *p4ha1a* genes were affected by cryopreservation with DMSO. Among the promoter regions affected by MeOH, one regulates *u2* gene. The other affected distal promoters were localized in the mitochondrial chromosome. These promoters are related to *cox2*, *cox3*, *nd3*, *nd4*, *nd4l*, *atp6* and *atp8* genes. The *pcdhb* and *slc26a2* genes had their first intron affected as well. We also found that among the distal promoters affected by glycerol, one was associated to *znf385d* gene and another one was associated to *osbpl10* gene. Other genes such as *ptprub* and *ndufs8a* were also affected at their introns by glycerol.

### Relationship between DNA methylation changes and cellular alterations

To explore the possible link between DNA methylation changes after cryopreservation and the corresponding cellular alterations of the same samples, we assessed whether there was a correlation between the number of DMCs and the fertilization rate. Indeed, fertilization ability is the cellular quality parameter that best reveals the overall functioning of the spermatozoa. No significant correlation (p > 0.05) was found between the 2 parameters. Because glycerol did not yield any fertilization, thus preventing the testing of some correlation, two other cellular parameters that are plasma membrane quality and mitochondrial activity were also tested. No correlation with DMCs number was found either, whatever the cryoprotectant. Surprisingly, a significant negative correlation was found between DMC number and motility after cryopreservation with DMSO (p = 0.04, not shown).

## DISCUSSION

Cryopreservation of rainbow trout sperm with 3 different cryoprotectants led to varying pattern of cellular alterations of thawed spermatozoa in our study. Hence, this validated our choice of the cryoprotectants, as they induced different cellular responses to cryopreservation. In addition, the variable response towards cryopreservation among fish males revealed an intrinsic biological variability. Such overall variability provided us with a set of samples that was suitable to study the effect of cryopreservation on the molecular parameter that is DNA methylation. The RRBS method used in this study yielded a limited number of cytosines in CpG sites, and this reduced set of data made possible the use of many biological replicates to explore the impact of sperm cryopreservation on DNA methylation. Our results demonstrated that cryopreservation had no straightforward impact on the global DNA methylation, although it yielded some DMCs between cryopreserved and fresh sperm. Presumed regions that would be more sensitive to cryopreservation were proposed, but few were shared between cryoprotectants.

Our study showed that the extent of the cellular cryoprotection provided by the different cryoprotectants had no related consequence on DNA methylation alteration. Indeed, using either an optimal (DMSO), non-optimal (MeOH) or the worst cryoprotectant (glycerol) did not affect the global DNA methylation in rainbow trout. This is at odd with some other species such as goldfish ^10^ and European eel ^9^, where the optimal cryoprotectant which is the MeOH did not alter global DNA methylation assessed by LUMA, while DMSO induced a hypomethylation. Therefore, our study revealed the stability of global DNA methylation towards the cryopreservation whatever the cryoprotectant used in rainbow trout. However, since we used RRBS technique, this allowed us to go further than global methylation and to assess DNA methylation at the nucleotide level.

We observed a limited number of DMCs after sperm cryopreservation, although a high number of paired cytosines was analyzed in each comparison. Besides, we were not able to find any DMR except 2 in glycerol. This further emphasizes how slight is the impact of cryopreservation on sperm DNA methylation in rainbow trout. This failure to detect more DMCs and DMRs is not due to the reduced representation of RRBS used here. Indeed, in a study dealing with rainbow trout physiological adaptation to its environment, 108 DMRs were found in spermatozoa between hatchery and natural origin rainbow trout using the same RRBS technique ^15^. We surmise that the impact of hatchery on DNA methylation was important enough to induce many DMCs leading to many DMRs. In our study, since the impact of sperm cryopreservation is tenuous, it yielded a restricted number of DMCs and as a consequence the DMCs were not dense enough to form DMRs. However, despite the limited number of DMCs found in our study, these DMCs are reproducible among the males since they were found in at least 6 males. This reveals the importance of having many biological replicates when assessing the impact of cryopreservation in order to validate the slight impact that can be observed.

Our analysis revealed that cryopreservation induced both hypomethylation and hypermethylation depending on the DMCs, so that no clear trend could be proposed as to the effect of cryopreservation. In addition, the status of the same DMC differs among the males, and the same male has DMCs with different status. Because of the quiescent state of sperm chromatin, the methylation changes were unlikely mediated by the enzymes responsible for methylation (the DNMTs) and demethylation (the TET pathway). Besides, the process of freezing-thawing of the spermatozoa is too fast for any enzyme being able to catalyze the methylation or demethylation reaction. The hypomethylations and hypermethylations observed in our study were more likely resulting from chemical impact on DNA methylation. Until now, it is still unclear how the cryoprotectants are able to modify the DNA methylation. DMSO is a chemically reactive molecule bearing 2 methyl groups, and it has been shown in an in vitro study that DNA can acquire methyl groups when incubated with DMSO and hydroxyl radicals ^16^. In the case of cryopreservation, the releasing of reactive oxygen species by damaged sperm mitochondria may trigger the formation of these hydroxyl radicals and induce this DMSO-associated methylation event. Regarding the demethylation event, it has been shown that 5-mC can be oxidized into 5-hmC in the presence of reactive oxygen species ^17,18^. The further transformation of 5-hmC into cytosine was shown to be made possible at high pH ^19^. It is known that pH can increase upon freezing of buffered solutions ^20^, and it cannot be excluded that a similar change is occurring locally within the cryopreserved spermatozoa. Therefore, demethylation may take place by chemically oxidizing 5-mC, and 5-hmC transformation into a cytosine may be linked to local pH changes during cryopreservation. The puzzling part in all these chemically-induced methylation changes is that both methylation and demethylation took place in the same sperm sample, and we have no clue as how such microreactors are made possible in the nucleus.

Despite the lack of DMRs in our study, we could identify potential sensitive regions in which several cytosines showed methylation changes in one or two cryoprotectants. Some of these regions are related to gene promoter regions that are instrumental to gene proper expression pattern. We focused our discussion on the genes affected by DMSO and MeOH, as glycerol cannot produce any offspring. Interestingly, the genes of promoter regions affected by cryopreservation with DMSO (*ebf2, p4ha1a*) are involved in specific processes of embryogenesis. In zebrafish, ebf2 (early B cell factor 2) is a transcription factor involved in primary neurogenesis in the embryo ^21^, where it stabilizes the commitment of the neuronal progenitors. The *p4ha1a* gene (prolyl 4-hydroxylase, alpha polypeptide I orthologue a) is an ortholog of *p4ha1* whose protein is involved in the regulation of early growth in bone tissues in bighead carp ^22^. Regarding the genes whose introns or promoters were affected after cryopreservation with MeOH, some had developmental functions as well (*pcdhb*, *slc26a2*, *u2*). While no function of *pcdhb* and *slc26a2* was described yet in fish, it has been shown in mice that *pcdhb* is involved in the regulation of neural development during the axonal assembly ^23^ and the axonal convergence into glomeruli in the olfactory bulb ^24^. Expression inhibition of slc26a2 (diastrophic dysplasia sulphate transporter) in mice is altering bone growth ^25^, and aberrant methylation of slc26a2 in human inhibits this gene expression and reduces its protein expression ^26^. Concerning *u2*, it has been shown in zebrafish that its protein belongs to the pre-mRNA Retention and Splicing (RES) complex that is fundamental during zebrafish embryogenesis ^27^. The other genes (*cox2*, *cox3*, *nd3*, *nd4*, *nd4l*, *atp6* and *atp8*) are involved in the oxidative phosphorylation (OXPHOS) reaction in fish as they belong to different mitochondria respiratory complexes ^28–30^. In light of the limited amount of data that was collected for each region, these affected genes represent preliminary clues about a slight impact of cryopreservation on DNA methylation, and provide a basis for further studies on important candidate genes. Recently, one study claimed that differentially methylated genes would be found after cryopreservation of Black Rockfish sperm ^31^, but the authors studied CpC dinucleotides instead of CpGs. These dubious methylation results on sites which are not supposed to be methylated in vertebrates unfortunately prevent any comparison with our present data. Indeed, our study was focused on the CpG sites, the only sites whose changes in methylation status are known to have functional consequences in vertebrates.

At this point, it is difficult to estimate whether the slight alterations observed in our study bear any consequence for the progeny. Among the thawed spermatozoa, there may be some whose DNA methylation was not affected by cryopreservation. If one of these spermatozoa is the one which fertilizes an oocyte, there will be no transmission of any DNA methylation alteration. Nevertheless, if a spermatozoon with an altered DNA methylation pattern did fertilize an oocyte, there may be a risk of transmitting this alteration to the progeny, although the extent of such transmission cannot be predicted. Indeed, although it is known that DNA strand breaks can be repaired by the maternal machinery in rainbow trout ^32^, nothing is known yet about DNA methylation repair. If the offspring is to inherit the altered DNA methylation pattern of the spermatozoa and that the affected genes are essential for embryo development, this should induce some increased embryonic mortalities. However, fertilization with cryopreserved sperm in rainbow trout was reported to be harmless regarding embryo development, once fertilization is triggered ^33,34^. It cannot be excluded however that non-lethal long-term effect could be transmitted. For example, rainbow trout embryos sired with cryopreserved spermatozoa had altered expression of several marker genes ^35^, and their stress resistance was modified at later stages ^36^. However, apart from the candidate gene analysis proposed above, an analysis of the long-term effect triggered by the tenuous changes observed in the present work seems a difficult task.

Our data revealed the absence of a link between alteration of DNA methylation and fertilization ability. The males that had the worst quality for a cellular quality parameter does not necessarily had the highest number of DMCs. This indicates that the damages which altered the spermatozoa functions to a great extent, such as cell dehydration and ice crystal formation ^37^, had little effect at the chromatin level. As a consequence, none of the cellular parameters used in this study can be used as markers of the risk of DNA methylation alterations. This lack of correlation can be explained by the general stability of the DNA methylation, validated by the low number of DMCs and the overall maintenance of the global DNA methylation level in all samples.

## MATERIAL AND METHODS

### Ethics statement

Rainbow trout males from the synthetic strain were kept in the experimental fish facilities ISC LPGP of INRAE (Agreement number D-35-238-6) with full approval for experimental fish rearing in strict line with French and European Union Directive 2010/63/EU for animal experimentation. The Institutional Animal Care and Use Ethical Committee in Rennes LPGP (Fish Physiology and Genomics Department) specifically approved this study (n° T-2020-37-CL). All fish were handled for gamete collection in strict accordance with the guidelines of the Institutional Animal Care and Use Ethical Committee in Rennes LPGP (Fish Physiology and Genomics Department). CL is accreted by the French Veterinary Authority for fish experimentation (n° 005239). The animal study is reported in accordance with ARRIVE guidelines (https://arriveguidelines.org) for animal research.

### Sperm collection and cryopreservation

Two years old mature males from the SYNTHETIC INRAE strain (average weight 1.2 kg) reared at the experimental fish farm UE PEIMA of INRAE were transferred to ISC LPGP fish facility. After 2 weeks in recirculated water tanks (12 °C, winter photoperiod), 12 fish were anaesthetized, and their sperm was collected by gentle striping and stored individually on ice. All sperm samples were treated individually (12 biological replicates in every analyses). They all had high motility and high membrane quality percentages (mean of the 12 males: 99.92 ± 0.2 and 98.05 ± 0.84 respectively). Fresh sperm concentration was adjusted to 6 × 10^9^ spermatozoa/mL in SFMM buffer (110 mM NaCl, 28.3 mM KCl, 1.1 mM MgSO_4_,7H_2_O, 1.8 mM CaCl_2_,2H_2_O, 10 mM Bicine, 10mM Hepes sodium salt, pH 8), and a fraction (50.10^6^ spermatozoa) was snap-frozen in liquid nitrogen prior to storage at - 80°C. This fraction was referred to as fresh sample, as it was not submitted to the whole cryopreservation process (no exposure to cryoprotectant, no slow freezing and thawing and no associated cellular constraints ^37^). The remaining sperm sample was split into 3 fractions to be cryopreserved with one of the three penetrating cryoprotectants dimethylsulfoxide (DMSO), methanol (MeOH) and glycerol (Figure 4).

**Figure 4:**
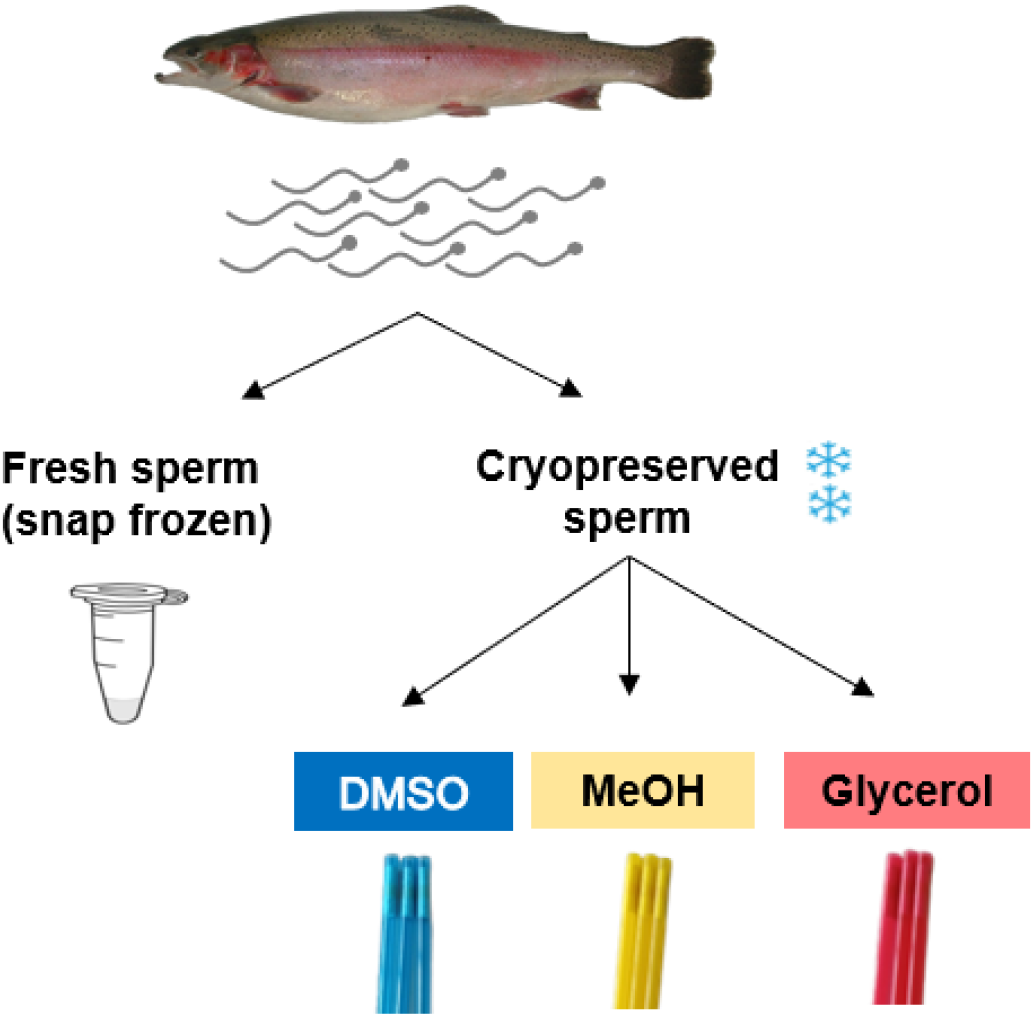
Experimental design. One sperm sample from each rainbow trout male (n=12) was split into 4 fractions: one fraction was snap frozen in liquid nitrogen and used as fresh control. The 3 other fractions were cryopreserved, each one with a different cryoprotectant: dimethyl sulfoxide (DMSO), methanol (MeOH), and glycerol.

Sperm cryopreservation was carried out with the FreezeFish kit (IMV Technologies ref 026913). One volume of sperm at 6 × 10^9^ cells/ml was mixed with one volume of each cryopreservation solution (buffer with sucrose and liposomes and one of the three penetrating cryoprotectant: DMSO, MeOH or glycerol). The final cryoprotectant concentration with the spermatozoa was 7.5 % (vol %). The mix of sperm suspension with each cryopreservation solution was loaded into 500 µL French straws (36 to 48 straws per male and 144 to 192 per condition). Straws were cryopreserved in the programmable freezer Micro DigitCool according to a fast-freezing curve (bovine curve, IMV Technology) and were then stored into liquid nitrogen until use.

When needed, the straws were thawed in a water bath at 37 °C for 10 s. Thawed sperm was then washed by dilution with SFMM (1 vol + 20 vol) and centrifugation at 200 g for 10 min at 4 °C. The pellet was resuspended at 100 × 10^6^ cells/mL in SFMM unless otherwise stated, and maintained on ice before use.

### Membrane quality

Membrane integrity of the spermatozoa was assessed from their ability to exclude the fluorescent probe propidium iodide. Fresh and washed frozen-thawed sperm samples diluted at 1 × 10^6^ cells/mL were incubated with propidium iodide (final concentration 12 µM) for 2 min and the percentage of unstained cells (cells with intact plasma membranes) was assessed by flow cytometry (MACSQuant Analyzer 10, Ref 130-096-343). Measurements were performed in duplicate on each sperm sample, using two different sperm suspensions in propidium iodide, within 1 h after thawing.

### Mitochondria quality

Mitochondrial membrane potential of the sperm samples was assessed with the JC-1 fluorescent probe using the kit “MitoProbe™ JC-1 Assay Kit for Flow Cytometry (M34152, Invitrogen by Thermo Fisher Scientific). Diluted sperm in SFMM was incubated for 15 min at 20 °C with JC-1 at a final concentration of 0.5 µM. JC-1 fluorescence was then measured by flow cytometry (MACSQuant Analyzer 10). The population with the highest red and green fluorescence was selected as the one whose spermatozoa had high quality mitochondria, and its representation was given as a percentage of the whole sperm population. Each sperm sample was measured in triplicates within 1 h after thawing.

### Motility

Thawed spermatozoa motility was assessed using a Computer-assisted Sperm Analysis (CASA) IVOS II using the software HTCasaII (IMV Technologies). The measurements were performed in triplicates for each sperm sample within 3 min post thawing on unwashed spermatozoa. For each replicate, 10 µL of thawed sperm (2.4 × 10^9^ spz/mL) were activated with 600 µL of Actifish (IMV Technologies), and a fraction (2.8 µL) was loaded within 10 s into the chamber of a Leja slide (Leja®, REF. 025107-025108). Motility was recorded from 10 s to 15 s post activation (8 movies of 30 pictures at a speed of 60 pictures/min), and spermatozoa with an average path velocity (VAP) ≥ 30 µm/s were counted as motile. Results are expressed as a percentage of motile spermatozoa.

### Fertilization ability

The fertilization test was conducted by adding 100 µL of thawed unwashed spermatozoa and 10 mL of Actifish (IMV Technologies) to approximately 150 oocytes (2 × 10^6^ spermatozoa/oocyte). As a control for the quality of the oocytes, 25 µL of fresh sperm and 10 mL of Actifish (IMV Technologies) were added to approximately 150 oocytes every 8 fertilization tests. All sperm samples (12 males × 3 cryopreservation conditions and the fresh controls) were tested for fertilization on the same egg batch obtained from the pool of 30 spawns (spring strain females from UE PEIMA, 3 years old). The spawns were collected one day before the experiment and transported over night at 4°C under oxygen to ISC LPGP. After fertilization, eggs were incubated in the dark at 10°C in recirculated water at the fish facility ISC LPGP. Fertilization rates were assessed from the development rates at eyes stage (about 20 days post fertilization). To do so, the eggs of each fertilization test were transferred into a petri dish and a picture was taken. The total number of eggs was counted with the macro VisEgg ^38^ on Fiji-ImageJ while the number of fertilized oocytes (embryos with developed eyes) was manually counted with the Fiji-ImageJ counter. Fertilization percentage was calculated by the following formula: (number of fertilized eggs/total number of eggs × 100). The normalized fertilization percentage used in the results correspond to: fertilization rate of each sample/average fertilization rate of fresh sperm × 100.

### DNA extraction

Thawed washed spermatozoa (4 × 10^6^ spermatozoa) were digested overnight at 42 °C in 800 µL TNES buffer (125 mM NaCl, 10 mM EDTA, 17 mM SDS, 4 M urea, 10 mM Tris-HCl, pH 8) and proteinase k (P6556; Sigma-Aldrich) at 75 µg/mL (final concentration). After cooling, 800 µL of phenol:chloroform:isoamyl alcohol (25:24:1 vol) were added to the 800 µL TNES with digested spermatozoa. After gentle shaking for 15 min at room temperature, the two phases were separated by centrifugation for 15 min at 8000 g at 4 °C. A fraction of 400 µL of the upper aqueous phase was mixed with 100 µL NaCl 5 M in water and 1 mL cold 100 % ethanol and incubated for 15 min at room temperature prior to centrifugation for 15 min at 12 000 g at 4 °C. The DNA pellet was washed with 100 µL 75 % ethanol and centrifuged for 15 min at 12 000 g at 4 °C. The new DNA pellet was left to air dry. It was then resuspended in RNAse (A7973; Promega) 100 µg/mL final concentration in nuclease-free water and incubated for 1 h at 37 °C. DNA concentration was measured with the Qubit3 Fluorometer – invitrogen by Thermo Fisher Scientific using the Qubit™dsDNA BR Assay kit (REF Q32853, invitrogen by Thermo Fisher Scientific) according to the manufacturer’s instructions.

For practical reasons, the same thawed sperm was used for membrane quality assessment and for DNA extraction. In parallel, the same thawed sperm but from different straws was used for motility and mitochondria quality assessment.

### RRBS library preparation, sequencing and data processing

DNA digestion with MspI, library preparation and bisulfite conversion of the 48 samples (12 males × 4 conditions) using Diagenode Premium RRBS kit were performed by the Integragen company (Evry, France). Paired-end sequencing (100 b read length) was carried out by the company on Illumina NovaSeq™ 6000, with S2 patterned flow cells. Because of some difficulties with the first sequencing round (too many reads with adapters, indicating too short inserts), some samples have been resequenced after some changes in fragment size selection. These yielded an average of 66.5 × 10^6^ paired end raw reads (133 × 10^6^ total raw reads). The sequencing quality of the raw data was checked with FastQC program (version: FastQC_v0.11.7) (https://www.bioinformatics.babraham.ac.uk/projects/fastqc/). Raw reads were trimmed using Trim Galore to remove adaptors sequences, the bases added during RRBS library preparation, and the bases with a phred score < 30 (https://github.com/FelixKrueger/TrimGalore version: TrimGalore-0.6.5). In order to exclude any risk of overestimation of DNA methylation due to a low bisulfite conversion (< 98 %), we calculated the bisulfite conversion rate and found a mean value 99.6 %.

Using bowtie2 aligner (version: bowtie2-2.3.5.1) available in Bismark bio-informatic software ^39^, trimmed reads were aligned to the reference genome (Omyk_1.0 genome version). Methylation status of every cytosine in CpG sites was extracted using “bismark_methylation_extractor”. These steps are available in the workflow which agrees to FAIR principles and is accessible online (https://forgemia.inra.fr/lpgp/methylome). All the samples had a median sequencing depth of the cytosines in CpG sites above 18 (table 1) indicating that the data per sample was sufficient. To visualize methylation levels distribution on the reference genome, BigWig files were obtained with bedGraphToBigWig ^40^ and methylation values were visualized with Integrative Genome Viewer ^41^. Because both 5-mC and 5-hmC are protected from bisulfite conversion into uracil ^42,43^, the cytosines claimed to be methylated in this RRBS study are in fact either methylated or hydroxy methylated, and the information about the proportion of 5-hmC is not available. However, because 5-hmC are mainly found at an early step of the demethylation process (reviewed by Kohli and colleagues ^44^), we consider that their merging with 5-mC did not induce any bias in the interpretation of the data.

**Table 1.** Summary of the sequencing data obtained in the RRBS analysis. Total raw reads number includes reads 1 and reads 2 (paired end sequencing). Reads with adapter is the number of reads whose insert is shorter than 100 bp. The % mapping efficiency = (number of reads that mapped only once on the trout genome/trimmed reads number) × 100. Sequencing depth is the number of times a cytosine has been sequenced, mapped to the reference genome and extracted as a cytosine in CpG sites. The % total genomic cytosines in CpG sites refers to (number of identified cytosines in CpG sites / number of cytosines in CpG sites in the reference genome -i.e. 70.6 × 10^6^-) × 100. Values are expressed as mean ± SD of the 12 males in each condition (Fresh, Cryopreserved in DMSO, MeOH, or Glycerol). Gly: glycerol.

| Condition | Total raw reads<br>x 10<sup>6</sup> | Reads with adapter<br>x 10<sup>6</sup><br>(% of total) | % mapping efficiency | Median sequencing depth of cytosines | Number of identified cytosines<br>x 10<sup>6</sup> | % total genomic cytosines in CpG sites |
| --- | --- | --- | --- | --- | --- | --- |
| **Fresh** | 137.0<br>± 34.4 | 72.0 (53.0)<br>± 16.5 (± 3.5) | 51.3<br>± 1.1 | 39.8 | 2.8<br>± 1.0 | 4.0<br>± 1.5 |
| **DMSO** | 109.9<br>± 15.4 | 57.4 (52.4)<br>± 7.2 (± 2.3) | 50.6<br>± 0.5 | 29.1 | 2.9<br>± 0.8 | 4.0<br>± 1.1 |
| **MeOH** | 151.4<br>± 45.6 | 77.9 (51.9)<br>± 21.2 (± 2.2) | 50.8<br>± 0.6 | 26.7 | 3.6<br>± 0.2 | 5.1<br>± 0.3 |
| **Gly** | 134.1<br>± 49.0 | 71.0 (53.1)<br>± 25.8 (± 3.3) | 50.7<br>± 0.9 | 36.1 | 2.7<br>± 1.0 | 3.8<br>± 1.4 |

**Table 2:**
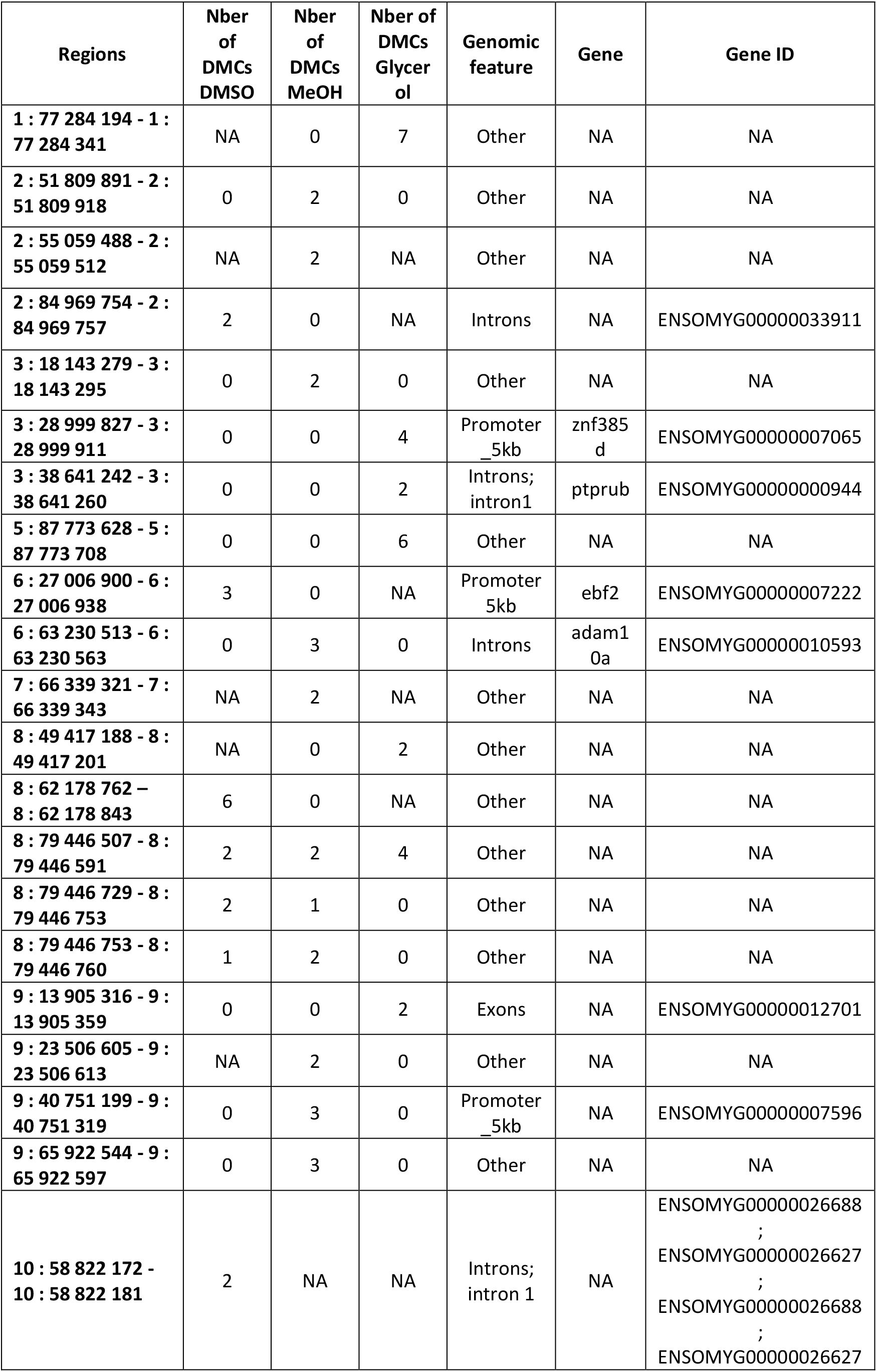
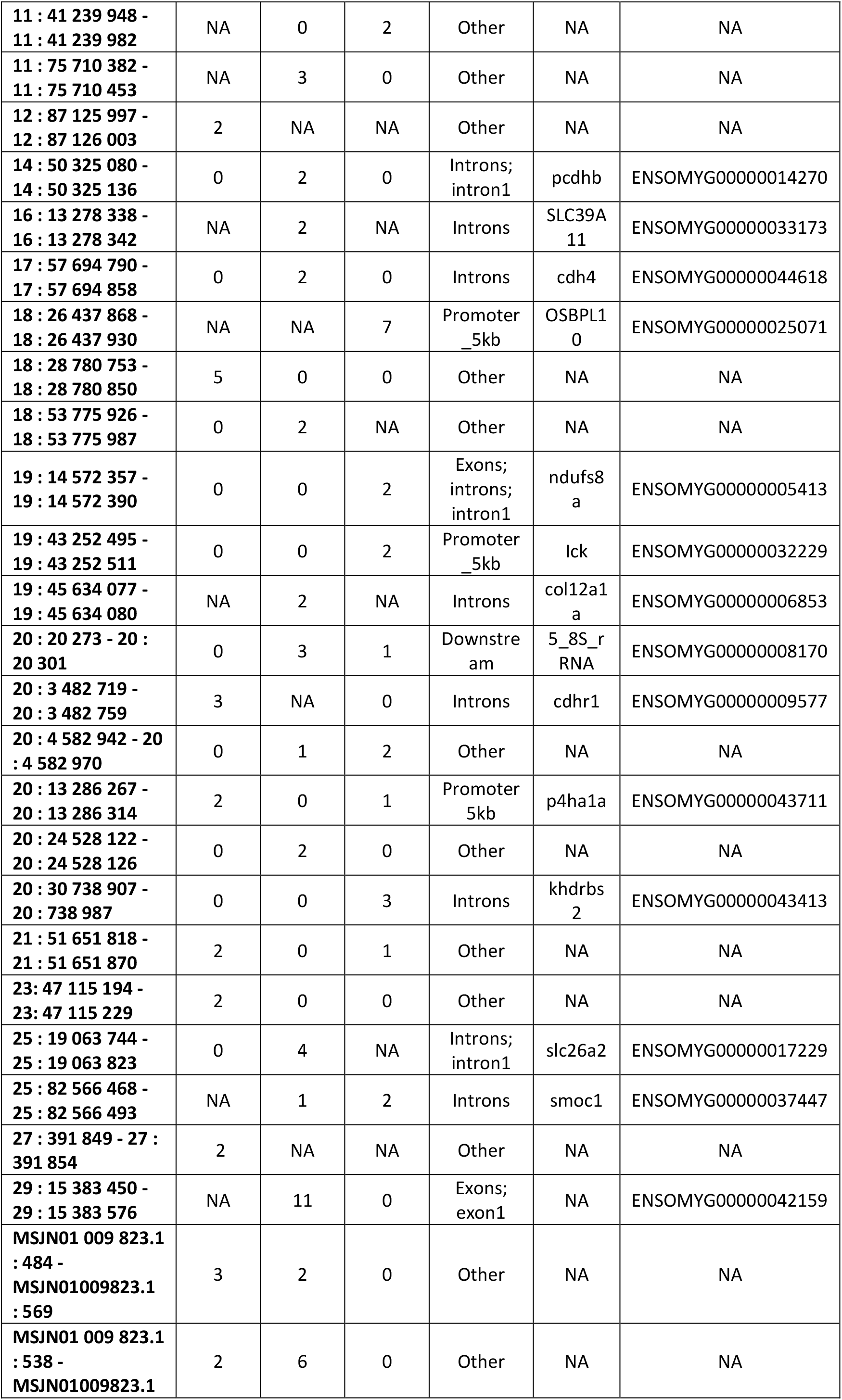
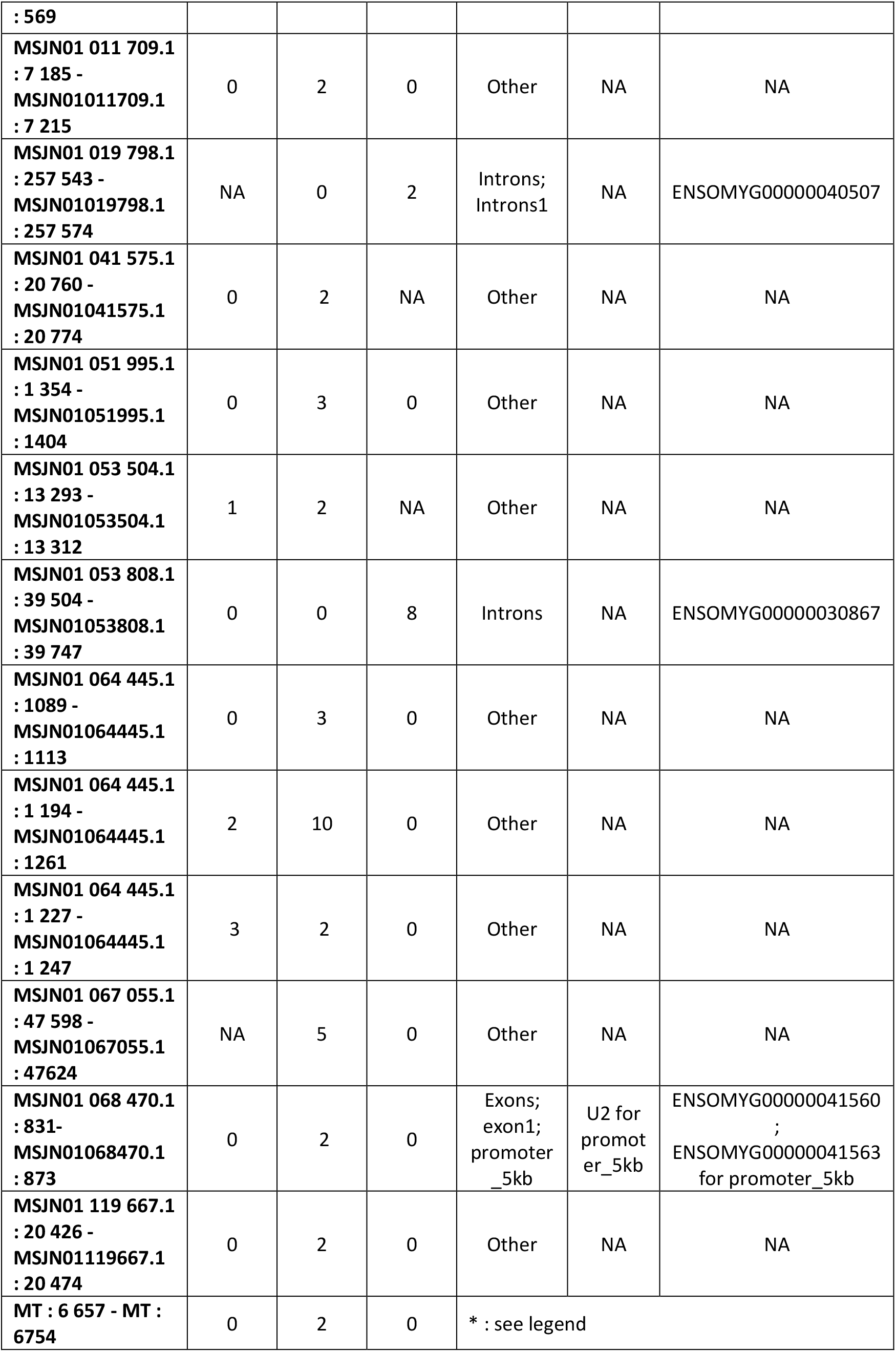
List of the potentially sensitive regions according to the cryoprotectant. The sequenced regions which displayed at least 2 DMCs (differentially methylated cytosines) between fresh and cryopreserved samples within 100 b (read size) were included in this list. The “Regions” column indicates the chromosome number and the region coordinates on the reference genome. Gene : name from the ensembl database (version 104) according to the reference genome (Omyk_1.0 genome version). Promoter 5 kb: region 5 kb upstream of the transcription starting site; Exon 1: first exon; Downstream: region 1 kb downstream of the transcription end site; Other: intergenic regions; NA: not available. *the 2 DMCs belonged to the exon1 of *cox1*, downstream of *nd2*, promoter 5 kb of *cox2*, promoter 5kb of *atp8*, promoter 5kb of *atp6*, promoter 5kb of *cox3*, promoter 5kb of *nd3*, promoter 5kb of *nd4l*, promoter 5kb of *nd4*.

For the identification of differentially methylated cytosines (DMCs), the paired design of DSS general experimental design from DSS ^45,46^ was used. For the DMCs detection, all cytosines with a sequencing depth N ≥ 1 were included in the analysis. Indeed, almost all cytosine sites are either completely methylated or unmethylated (Figure 3b), meaning that averaging the methylation status of several cytosines at the same site is not absolutely mandatory. In addition, DSS was also used to detect the differentially methylated regions (DMRs). The criteria used to determine the DMRs were to have a sliding sequence of at least 50 bases with ≥ 5 CpGs with a minimum of 75 % DMCs (with p-value < 0.001). Ven diagram was created using Jvenn ^47^, a web-based tool. The position of the DMCs within genomic features and the identity of the genes corresponding to regions potentially sensitive to cryopreservation was obtained by annotating DMCs with GenomeFeatures ^48^and ensembl v104.1 annotation.

## Acknowledgment

The authors acknowledge the skillful involvement of the staff from the ISC LPGP and UE PEIMA at INRAE for animal rearing and care. The authors gratefully acknowledge the help of Jerome Bugeon who devised the automatic counting of eggs. MEK is recipient of a PhD fellowship ARED from region Bretagne and INRAE PHASE (EPICRYSP 2020-2023). This work was funded by the French CRB Anim project ANR-11-INBS-0003 and the European FEAMP BIOGERM measure 47 (Innovation Aquaculture).

## Author contribution

MEK organized and carried out the study, analyzed and interpreted the data, and drafted the manuscript. AB taught and supervised the bioinformatic work of MEK, and contributed to the bio-informatic analyses. AL and CL conceived, designed and supervised the study. Additionally, CL supervised the writing of the manuscript and AL provided her knowledge on genomic DNA methylation in trout. All authors approved the final version of the manuscript.

## Data availability

The data for this study has been deposited in the European Nucleotide Archive (ENA) at EMBL-EBI under accession number PRJEB60579 (https://www.ebi.ac.uk/ena/browser/view/ PRJEB60579).”

## Competing Interests

The authors declare no competing interests.

## Notes

### Competing Interest Statement

The authors have declared no competing interest.

